## Supplementary Information for "Structural and Dynamic Impacts of Single-atom Disruptions to Guide RNA Interactions within the Recognition Lobe of *Geobacillus stearothermophilus* Cas9"

**Figure S1.** Amino acid sequence and structural alignments of *Geo* vs *Spy*Rec domains.
**Figure S2.** Assigned backbone $^{1}H$-$^{15}N$ TROSY HSQC spectra of *Geo*Rec1, *Geo*Rec2, and *Geo*Rec.
**Figure S3.** Overlay of $^{1}H$-$^{15}N$ TROSY HSQC and $^{1}H$-$^{13}CH_3$ ILV-methyl spectra of *Geo*Rec1, *Geo*Rec2, and *Geo*Rec.
**Figure S4.** Temperature-dependent CD spectra of *Geo*Rec1, *Geo*Rec2, and *Geo*Rec.
**Figure S5.** Structural perturbations of K267E and R332A *Geo*Rec2 identified by NMR.
**Figure S6.** Raw and temperature-dependent CD spectra of WT, K267E, and R332A *Geo*Rec2.
**Figure S7.** Summary of $R_1$, $R_2$, and $^{1}H$-[$^{15}N$] NOE NMR relaxation parameters, and $S^2$ for K267E and R332A *Geo*Rec2.
**Figure S8.** CPMG relaxation dispersion curves for WT *Geo*Rec2
**Figure S9.** CPMG relaxation dispersion curves for K267E *Geo*Rec2
**Figure S10.** CPMG relaxation dispersion curves for R332A *Geo*Rec2
**Figure S11.** Optimization of RNA sequences for titration into *Geo*Rec via NMR.
**Figure S12.** Structures of full-length and truncated *Geo*Cas9 RNAs
**Figure S13.** Summary of RNA NMR titrations using WT, K267E, and R332A *Geo*Rec2 subdomains.
**Figure S14.** Summary of RNA-induced change in NMR $\Delta\delta$ of WT, K267E, and R332A *Geo*Rec.
**Figure S15**. Stabilizing effect of gRNA on *Geo*Cas9 measured via CD.
**Figure S16.** MD simulations of WT, K267E, R332A, and K267E/R332A *Geo*Cas9 and i*Geo*Cas9
**Figure S17.** Summary of temperature-dependent *in vitro* cleavage of WT, K267E, and R332A *Geo*Cas9.
**Figure S18.** Summary of *in vitro* specificity assay for WT, K267E, R332A, and K267E/R332A *Geo*Cas9.
**Table S1.** Profiles fit to a global $k_{ex}$, determined from $^{1}H$-$^{15}N$ CPMG relaxation dispersion of WT, K267E, and R332A *Geo*Rec2.
**Table S2.** Nucleic acid sequences used in *in vitro* specificity assay for *Geo*Cas9.
**Table S3.** Nucleic acid sequences used in *in-vitro* specificity assay for *Sp*Cas9.

**Table S4.** Guide RNA sequences used in MST and NMR studies of *Geo*Cas9.

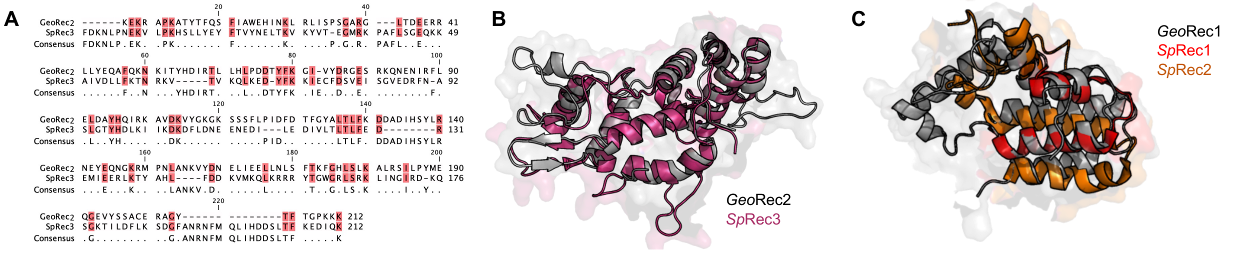


**Figure S1. (A)** Sequence alignment of *Geo*Rec2 and *Spy*Rec3. Conserved residues are highlighted pink and are listed as consensus. **(B)** Overlay of the *Geo*Rec2 X-ray crystal structure (gray) with *Spy*Rec3 from full-length *Spy*Cas9 (pink, PDB: 4UN3). **(C)** Overlay of the *Geo*Rec1 structure derived from full-length *Geo*Cas9 modeled with Alphafold2 (gray) with the homologous portion of *Spy*Rec1and *Spy*Rec2 from *Spy*Cas9 (PDB: 4UN3) in red and orange, respectively.

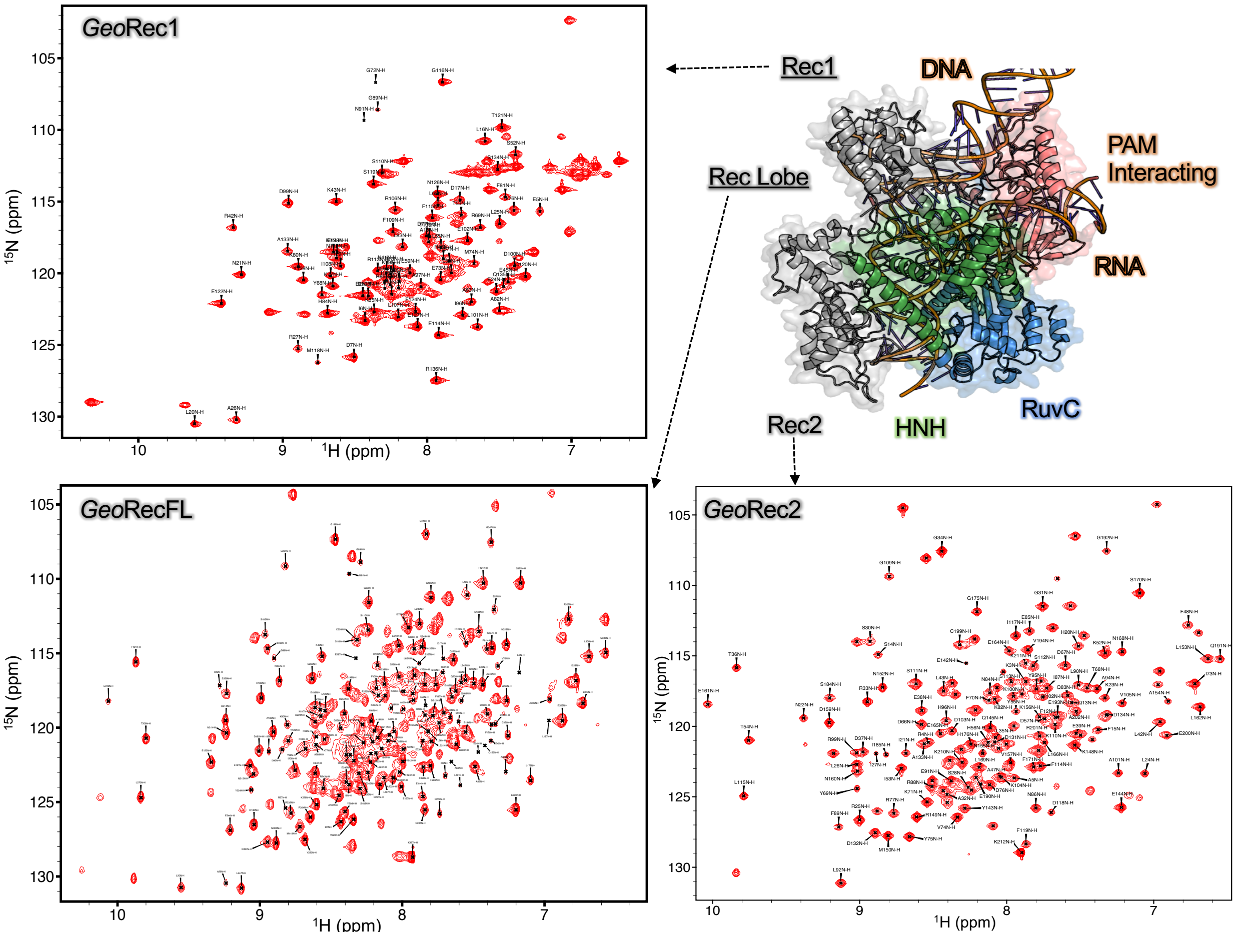


**Figure S2.** Assigned $^{1}H$-$^{15}N$ TROSY HSQC spectrum of *Geo*Rec1 (top left), *Geo*Rec (bottom left), and *Geo*Rec2 (bottom right).

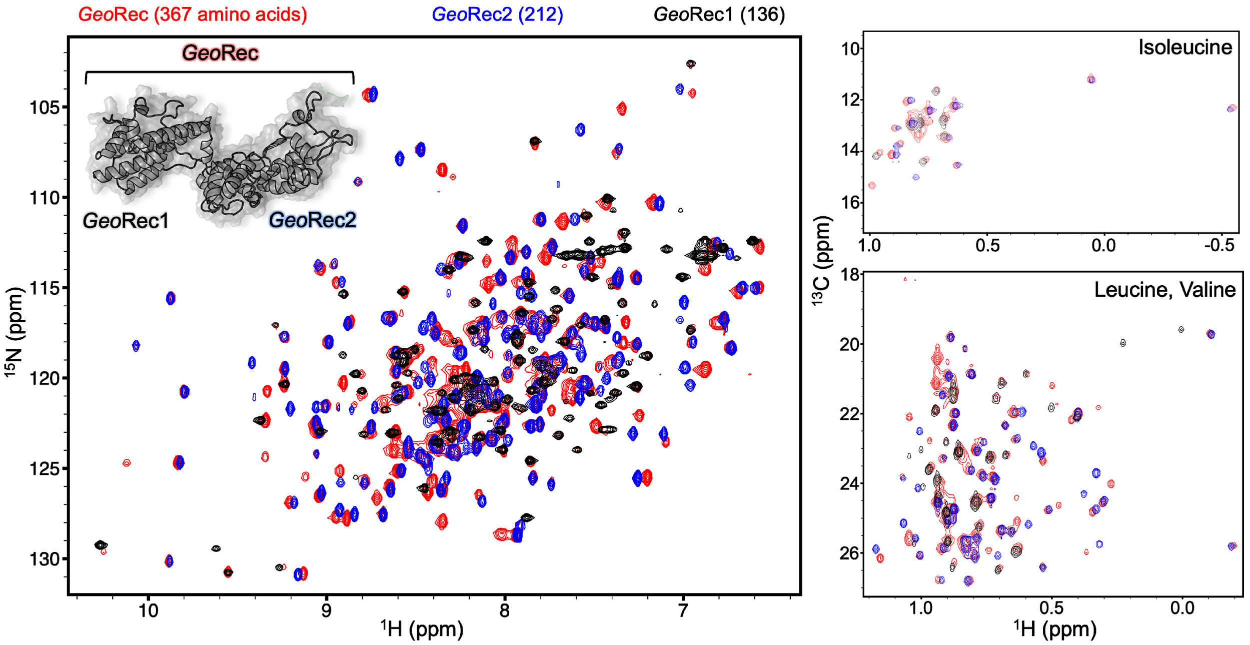


**Figure S3.** The dumbbell shape of *Geo*Rec is composed of the *Geo*Rec1 and *Geo*Rec2 subdomains. Overlay of $^{1}H$-$^{15}N$ TROSY HSQC spectra of *Geo*Rec (red), *Geo*Rec1 (black), and *Geo*Rec2 (blue) collected at 850 MHz shows that spectra of isolated *Geo*Rec subdomains overlay nicely with the spectrum of *Geo*Rec. The number of amino acids that make up each construct are indicated above the spectrum. Overlaid $^{1}H$-$^{13}CH_3$ ILV-methyl spectra of *Geo*Rec (red), *Geo*Rec1 (black), and *Geo*Rec2 (blue), show a similar property of the subdomains.

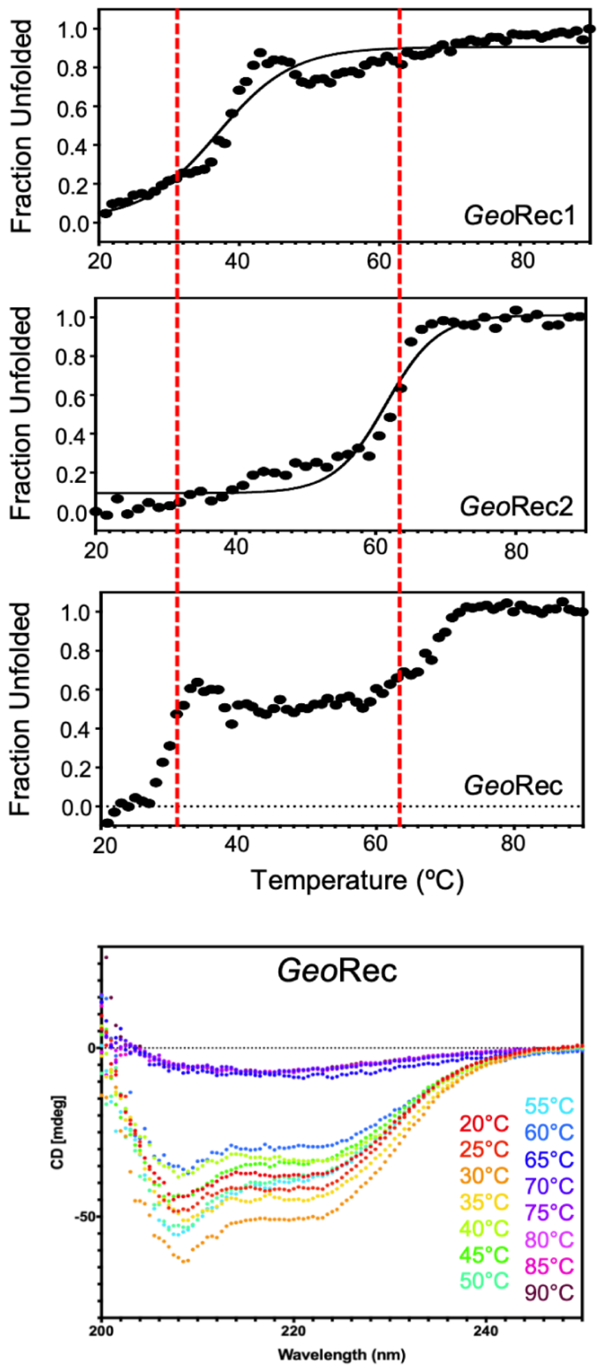


**Figure S4.** Temperature-dependent CD unfolding profiles of *Geo*Rec1, *Geo*Rec2, and *Geo*Rec reveal that the unfolding profile of the individual subdomains are conserved within that of *Geo*Rec. CD spectra spanning 200 – 250 nm at increasing temperatures (20-90 °C, bottom) show a gradual loss of *Geo*Rec secondary structure to ~60 °C, followed by an abrupt and complete unfolding at 65 °C, beyond the $T_m$ of *Geo*Rec2. The $T_m$ of *Geo*Rec1 and 2 were determined by fitting the CD data as described in the Materials and Methods. The red dashed lines indicate the $T_m$ values of *Geo*Rec1 and *Geo*Rec2, to guide the interpretation of the full-length *Geo*Rec unfolding profile.

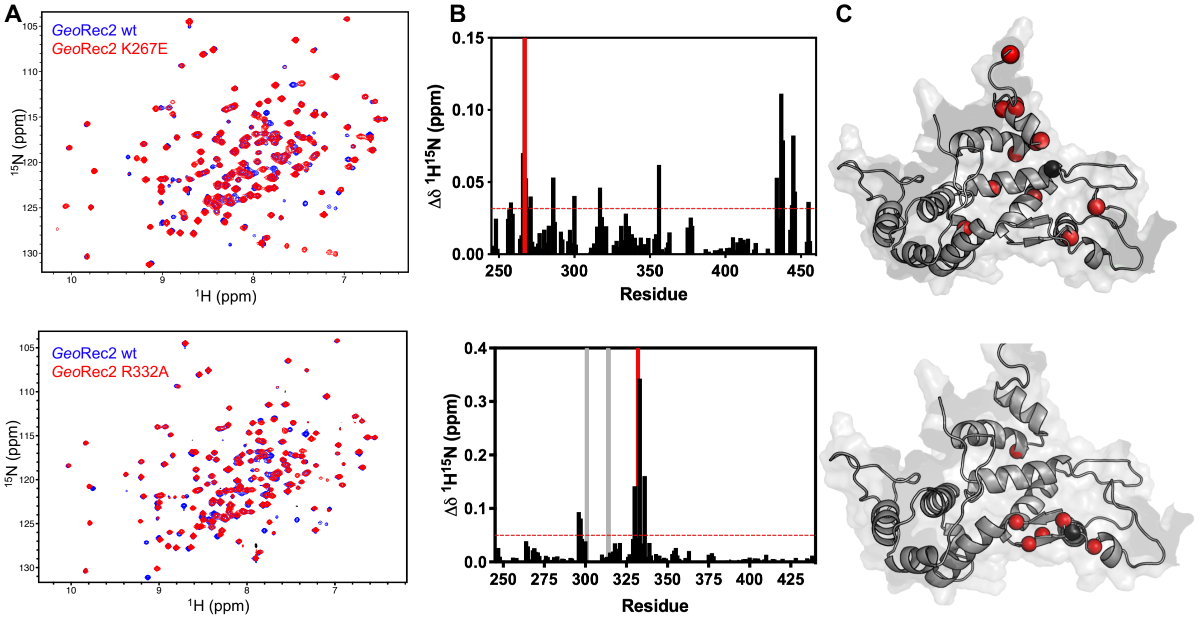


**Figure S5. (A)** 600 MHz $^{1}H$-$^{15}N$ TROSY HSQC spectra of WT *Geo*Rec2 (blue) overlaid with K267E *Geo*Rec2 (red, top) and R332A *Geo*Rec2 (red, bottom). **(B)** Chemical shift perturbations caused by K267E (top) or R332A (bottom) mutations are plotted for each residue. Resonances broadened beyond detection are marked with gray bars. The mutation site is marked with a red bar. **(C)** Chemical shift perturbations >1.5σ of the 10% trimmed mean of all shifts are shown as red spheres on the *Geo*Rec2. The mutation site is shown as black sphere.

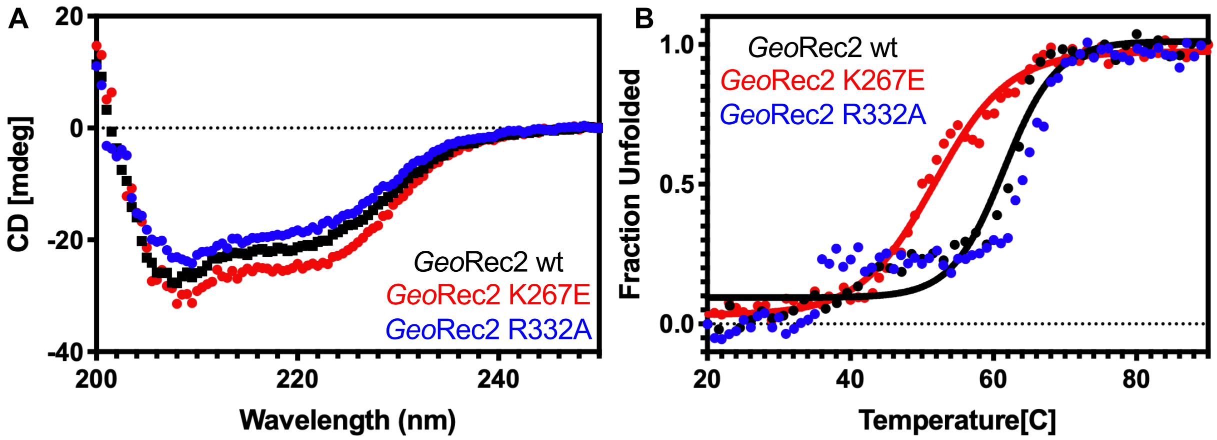


**Figure S6. (A)** Raw CD spectra of WT *Geo*Rec2 (black), K267E *Geo*Rec2 (red), and R332A *Geo*Rec2 (blue). **(B)** Temperature-dependent CD spectra reveal that K267E *Geo*Rec2 unfolds at a lower temperature (~55 °C, red) than WT *Geo*Rec2 (~62 °C, black). R332A *Geo*Rec2 undergoes a smaller unfolding event around 40 °C before completely unfolding ~62 °C (blue). The data were fit as described in the Materials and Methods.

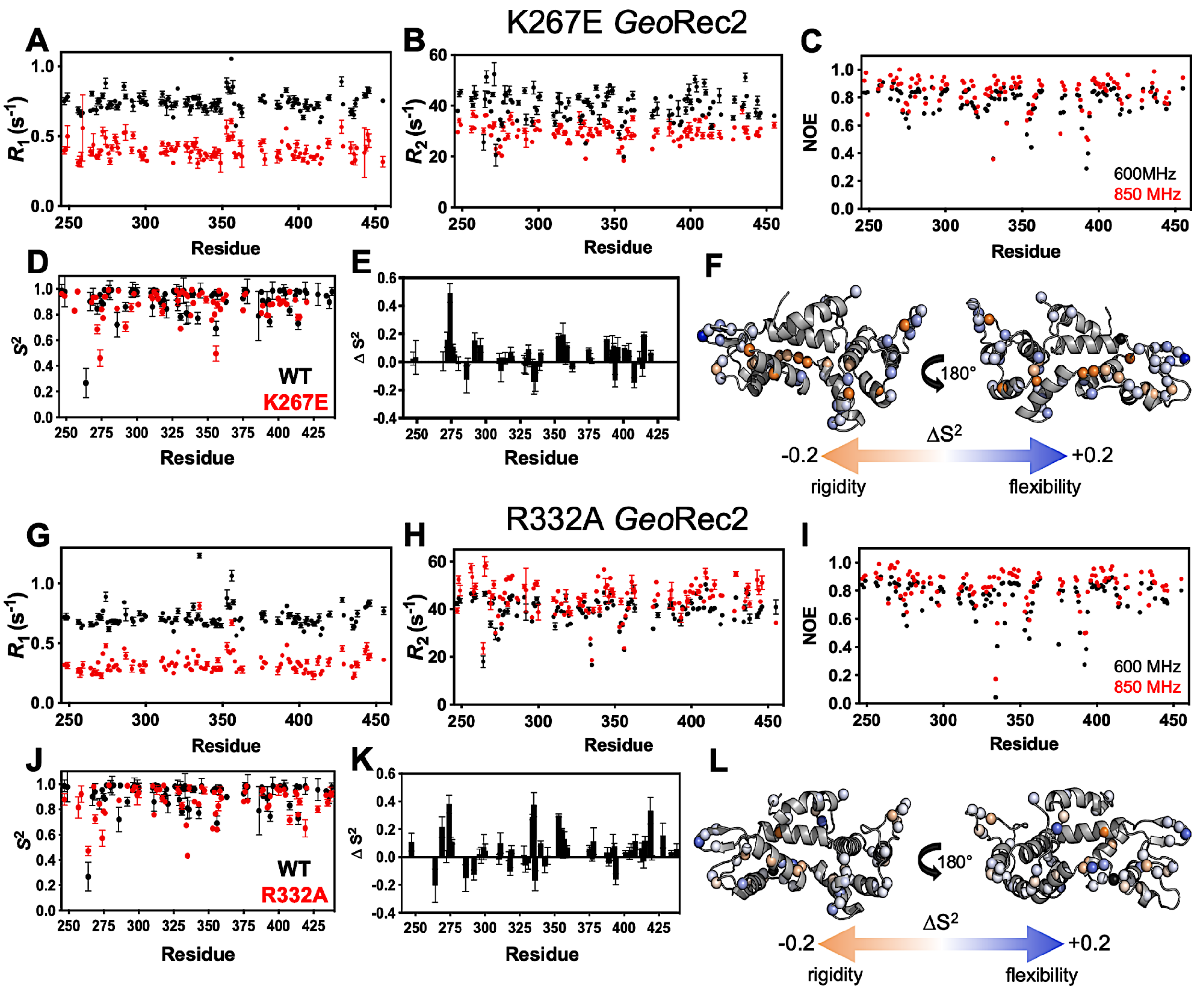


**Figure S7.** $R_1$ **(A)**, $R_2$ **(B)**, and $^1$H-[$^{15}$N] NOE **(C)** values for K267E *Geo*Rec2 measured by NMR at 600 MHz (black) and 850 MHz (red). **(D)** Order parameters ($S^2$) for K267E *Geo*Rec2 determined from Model-free analysis of $R_1$, $R_2$, and $^1$H-[$^{15}$N] NOE measurements (red) overlaid with $S^2$ values of WT *Geo*Rec2 (black). $\Delta S^2$ (plotted as WT – mutant) for K267E *Geo*Rec2 **(E)**, where a negative or positive values correspond to suppressed or heightened ps-ns flexibility of that site, respectively. K267E $\Delta S^2$ values are mapped onto the *Geo*Rec2 X-ray crystal structure in **(F)**. $R_1$ **(G)**, $R_2$ **(H)**, and $^1$H-[$^{15}$N] NOE **(I)** values for R332A *Geo*Rec2 measured by NMR at 600 MHz (black) and 850 MHz (red). **(J)** Order parameters ($S^2$) for R332A *Geo*Rec2 (red) overlaid with $S^2$ values of WT *Geo*Rec2 (black). $\Delta S^2$ (plotted as WT – mutant) for R332A *Geo*Rec2 **(K)**band R332A $\Delta S^2$ values are mapped onto the *Geo*Rec2 X-ray crystal structure in **(L)**.

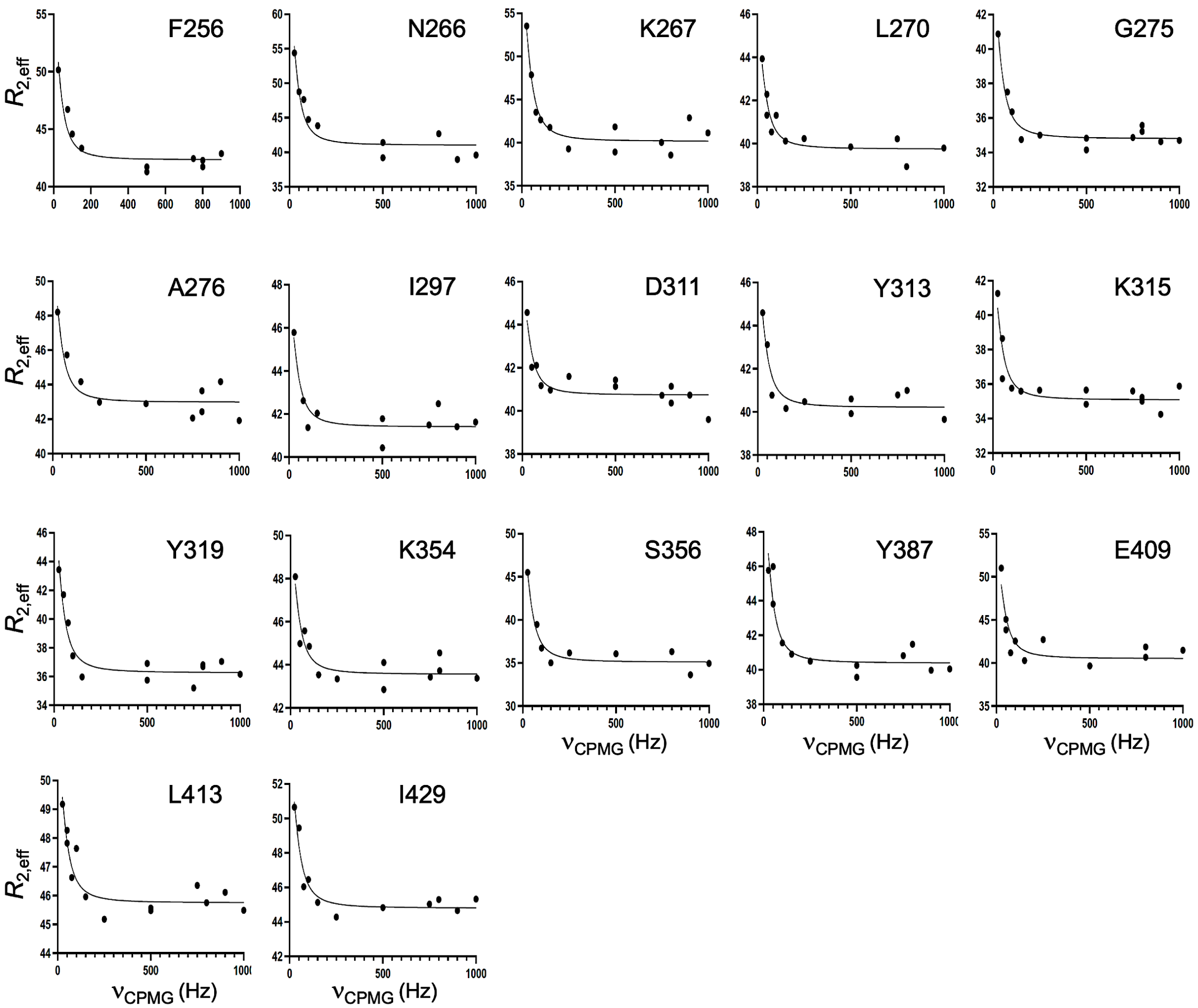


**Figure S8.** CPMG relaxation dispersion curves collected at 25 °C and 600 MHz for WT *Geo*Rec2. A global fit of all dispersion curves was determined to be superior based on the Akaike Information Criterion.

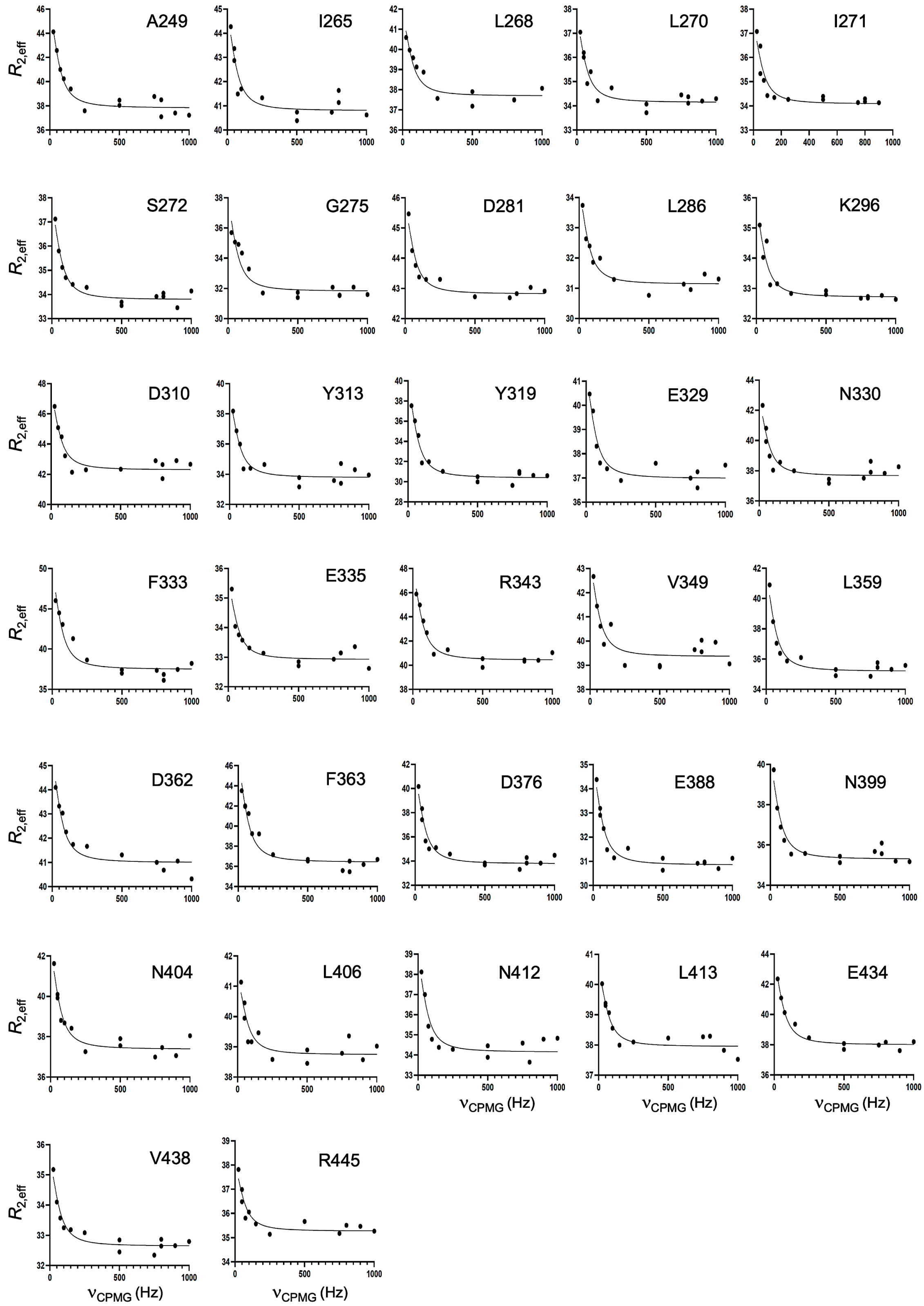


**Figure S9.** CPMG relaxation dispersion curves collected at 25 °C and 600 MHz for K267E *Geo*Rec2. A global fit of all dispersion curves was determined to be superior based on the Akaike Information Criterion.

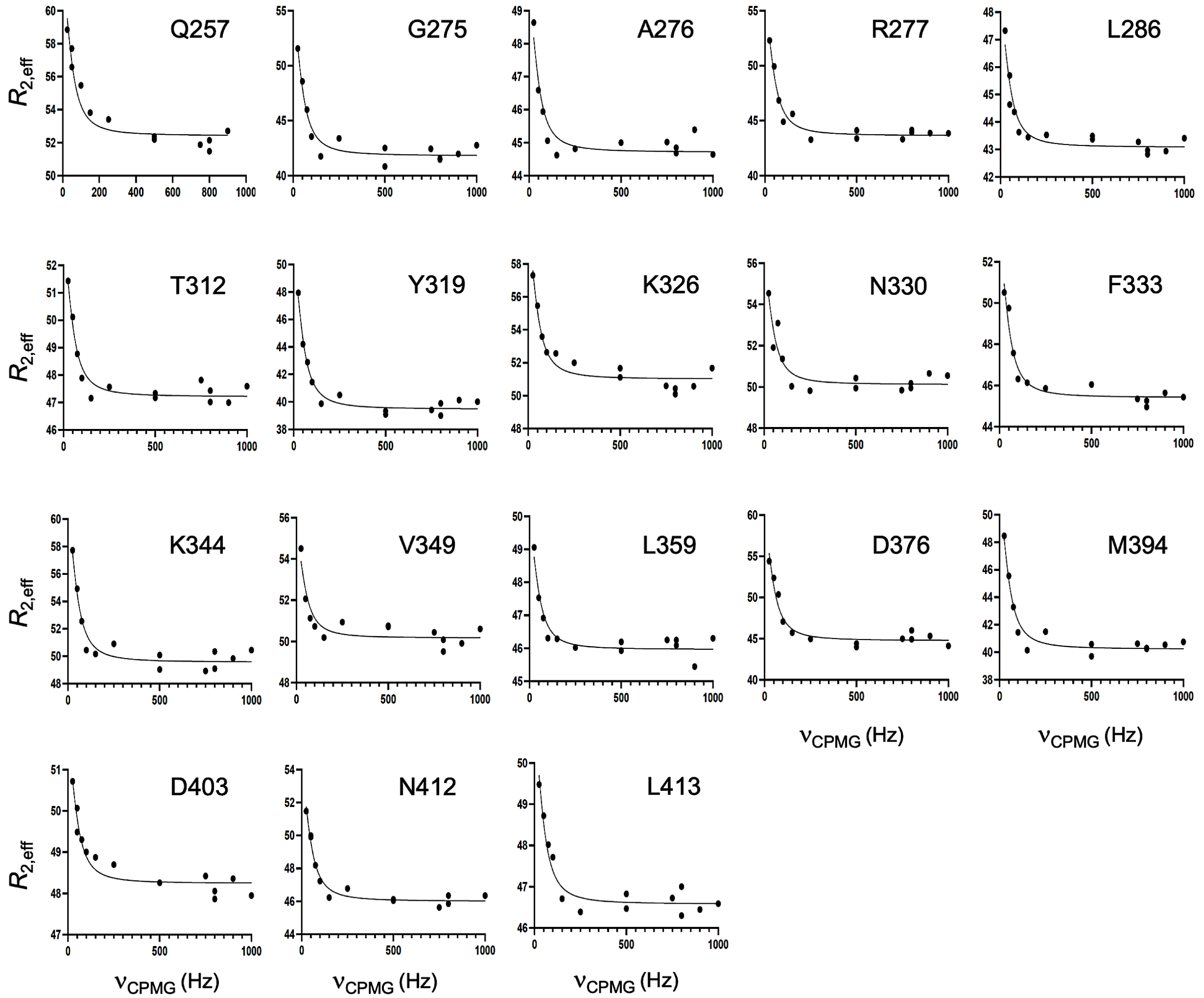


**Figure S10.** CPMG relaxation dispersion curves collected at 25 °C and 600 MHz for R332A *Geo*Rec2. A global fit of all dispersion curves was determined to be superior based on the Akaike Information Criterion.

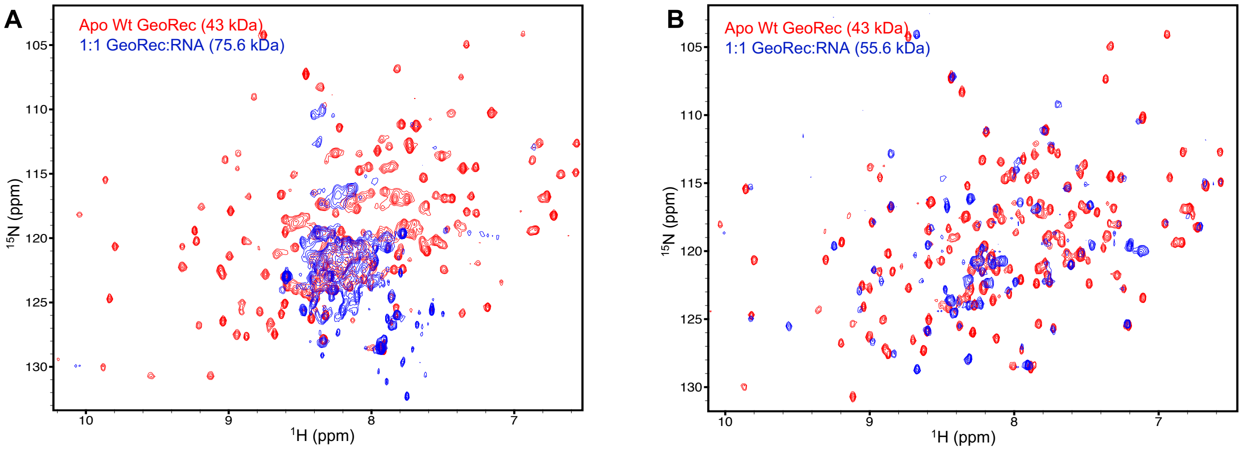


RNA used: 101nt (~32.5 kDa)
UUGCACCUACCUUCUGGAUGUAC*GUCAUAGUU*CCCCUGAGAA
AUCAGGGUUACUAUGAUAAGGGCUUUCUGCCUAAGGCAGACU
GACCCGCGGCGUUGGGG

RNA used: 39 nt (~12.6kDa)
UUGCACCUACCUUCUGGAUGUAC*GUCAUAGUU*CCCCUGA

Standard sgRNA: 141nt
UUGCACCUACCUUCUGGAUGUAC*GUCAUAGUU*CCCCUGAGAAAUCAGGGUUACUAUGAUAAGGGCUUUCUGC
CUAAGGCAGACUGACCCGCGGCGUUGGGGAUCGCCUGUCGCCCGCUUUUGGCGGGCAUUCCCCAUCCUU

**Figure S11.** 850 MHz $^{1}H$-$^{15}N$ TROSY HSQC NMR spectra of apo-*Geo*Rec (red) and *Geo*Rec in complex with either a 101-nt **(A)** or 39-nt portion of the full gRNA **(B)** at a 1:1 molar ratio (blue). The RNA sequence used in each NMR binding experiment is shown below each spectrum, along with the full-length gRNA sequence.

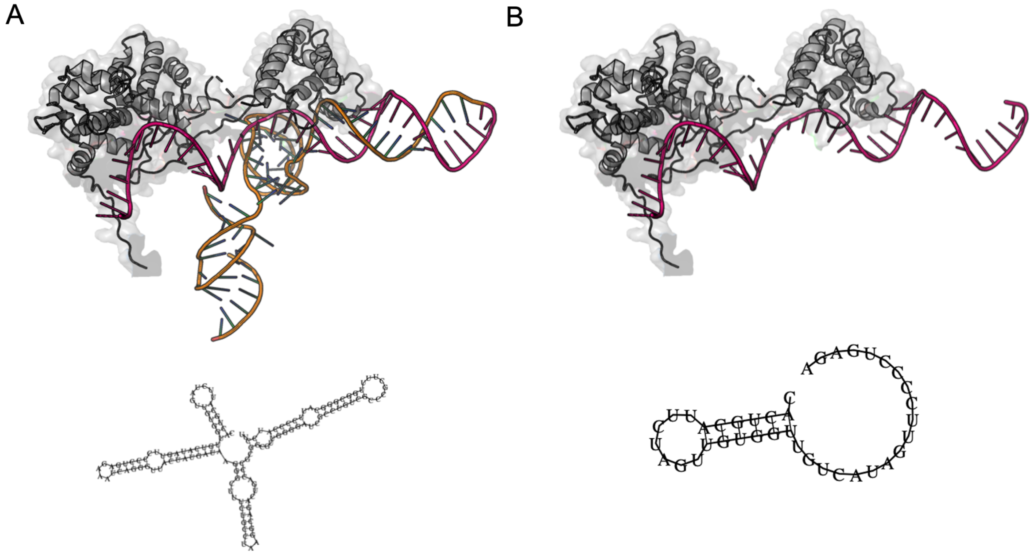


**Figure S12.** Structures of full-length **(A)** and truncated **(B)** *Geo*Cas9 gRNAs used in this work, based on the cryo-EM structure PDB: 8UZA. The 2D structure cartoons below each representation were predicted from sequence by the RNAfold web server (http://rna.tbi.univie.ac.at/cgi-bin/RNAWebSuite/RNAfold.cgi) maintained by the Institute for Theoretical Chemistry, University of Vienna.

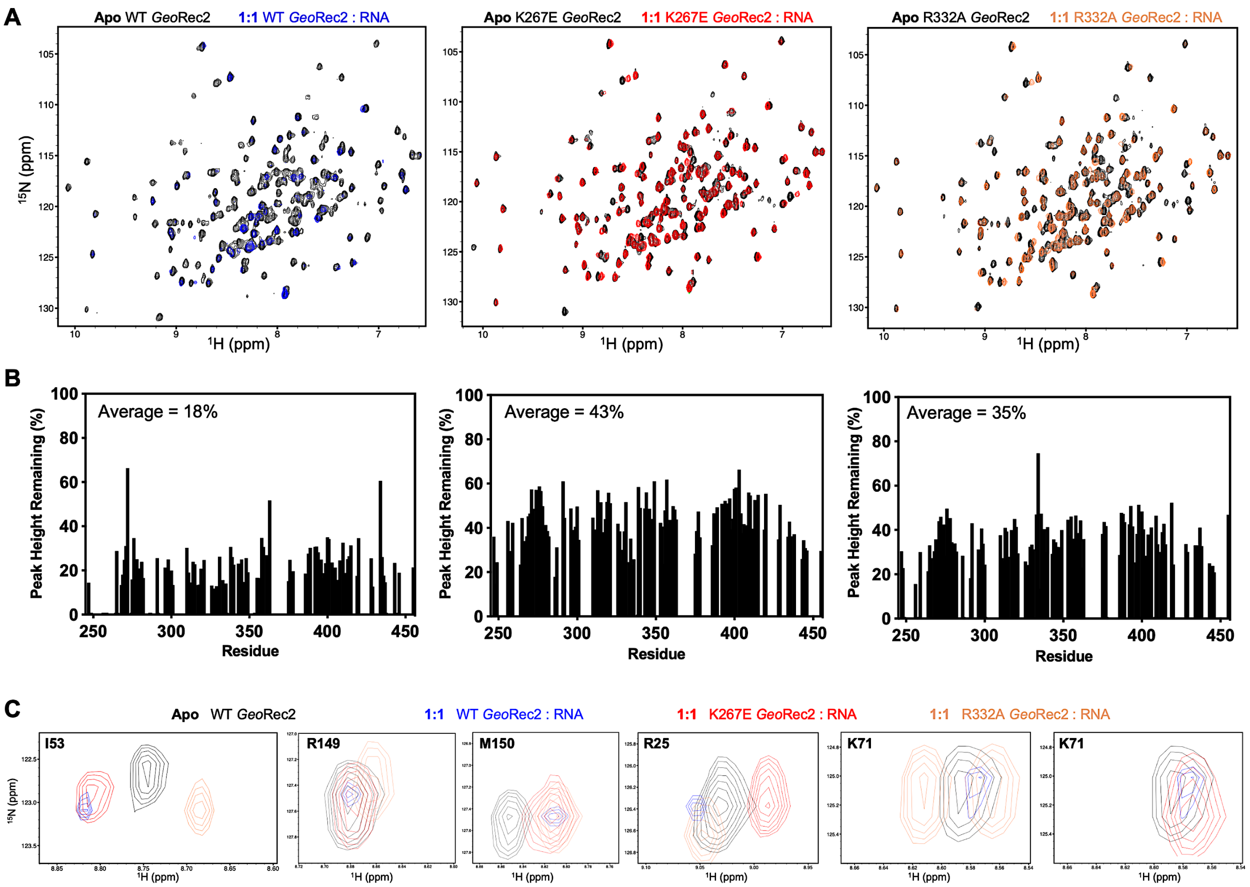


**Figure S13. (A)** $^{1}H^{15}N$ TROSY HSQC NMR spectral overlays of apo (black) and RNA-bound WT (blue), K267E (red), and R332A (orange) *Geo*Rec2. **(B)** Per-residue residual peak heights after addition of equimolar RNA to WT (left), K267E (center), and R332A (right) *Geo*Rec2. **(C)** NMR snapshots of apo WT *Geo*Rec2 (black) and gRNA-bound WT (blue), K267E (red), and R332A (orange) *Geo*Rec2, highlighting unique RNA-induced structural perturbations of each variant.

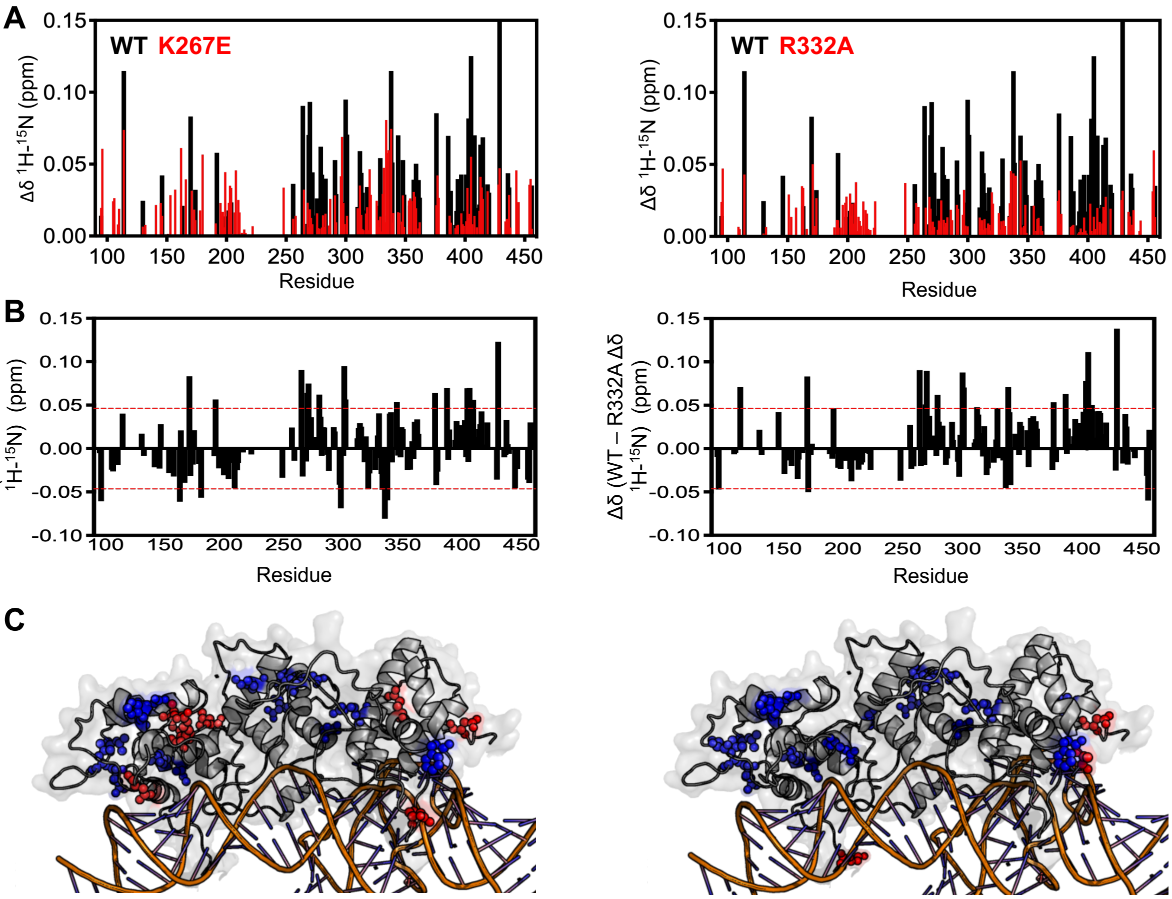


**Figure S14. (A)** RNA-induced NMR chemical shift perturbations to WT *Geo*Rec (black) overlaid with those of K267E (left) and R332A (right) *Geo*Rec, each in red. Similar plots are also found in Figure 1D and 4B of the main text. **(B)** The difference in RNA-induced chemical shift perturbation (Δδ WT – mutant) comparing WT and K267E *Geo*Rec (left), and WT and R332A *Geo*Rec (right). The red dashed lines indicate 1σ above and below the 10% trimmed mean of all shifts. **(C)** Chemical shift perturbations +1.0σ above the trimmed mean of all data are mapped onto *Geo*Rec as blue spheres, while perturbations -1.0σ are mapped onto *Geo*Rec as red spheres, representing key residues influencing RNA interactions.

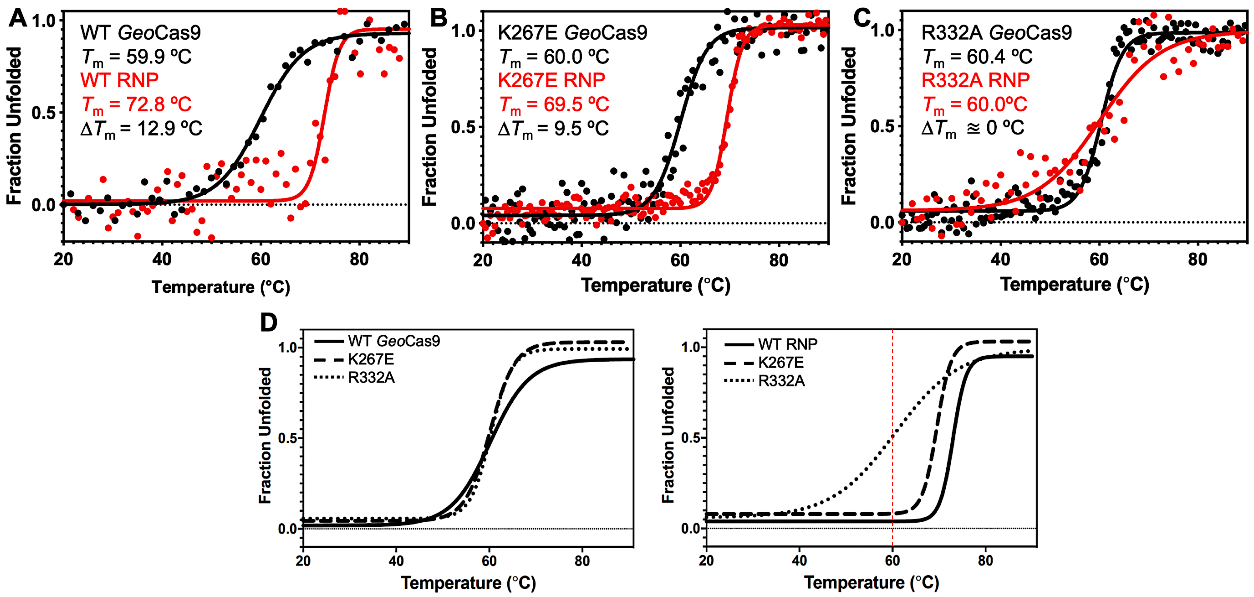


**Figure S15.** Stabilizing effect of gRNA on *Geo*Cas9. **(A)** CD spectroscopic unfolding profiles of WT *Geo*Cas9 (black) reveal a marked gRNA-dependent stabilization (red). The $T_m$ of each state is inset, as is the $\Delta T_m$. The same CD profiles for K267E *Geo*Cas9 **(B)** highlights a weaker stabilization and $\Delta T_m$ upon gRNA binding, while those of R332A *Geo*Cas9 **(C)** show virtually no difference in $T_m$ between apo and RNP samples. **(D)** Fitted denaturation profile overlays of apo WT (black line), K267E (dashed line), and R332A (dotted line) *Geo*Cas9 (left) and RNP complexes (141nt gRNA, right). The red dashed line denotes the $T_m$ of the apo proteins, which are nearly identical. All data were fit as described in the Materials and Methods.

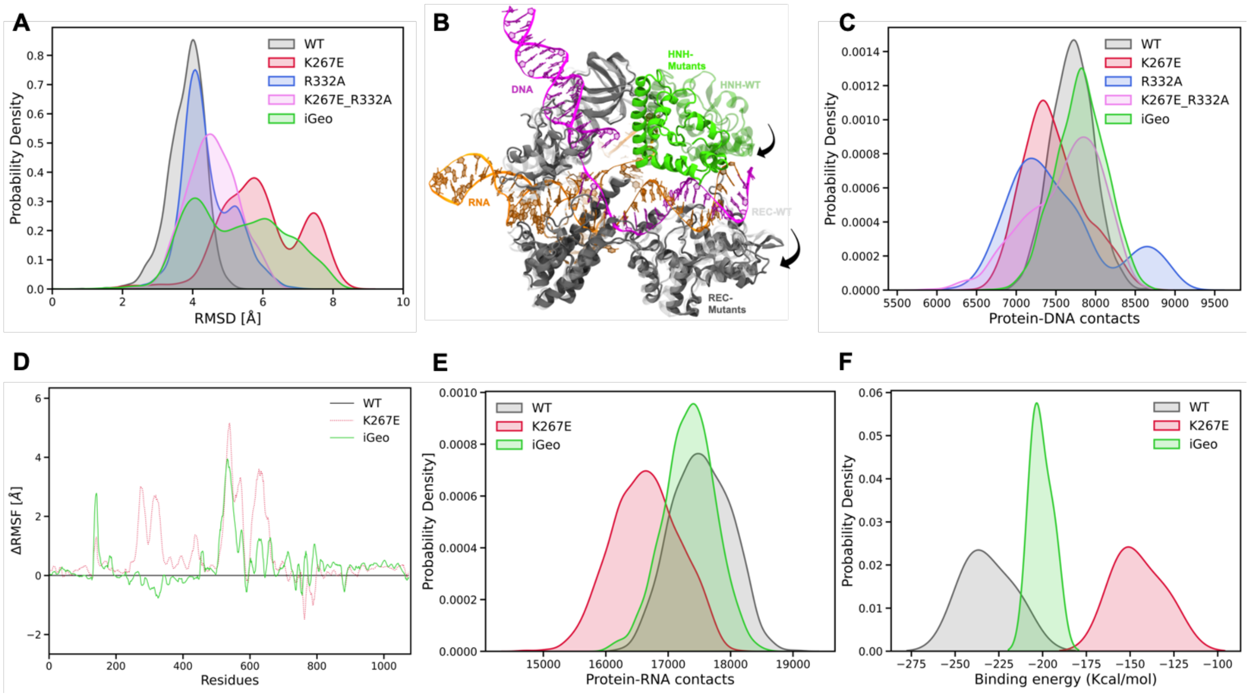


**Figure S16.** Analysis of MD simulations. **(A)** Root-mean-square deviation (RMSD) distribution of WT (gray), K267E (red), R332A (blue), K267E/R332A double mutant (pink) and i*Geo*Cas9 (green). **(B)** Conformational changes in the HNH and Rec domains of the mutant systems (black arrows), compared to WT *Geo*Cas9. **(C)** Distribution of protein-DNA contacts for WT and mutants computed over the 6 μs ensemble. **(D)** Differential RMSD (ΔRMSF) of the protein residues computed between the WT *Geo*Cas9 and K267E (red) and i*Geo*Cas9 (green). **(E)** Distribution of protein-RNA contacts for WT, K267E and i*Geo*Cas9 computed over the 6 μs ensemble. **(F)** Comparison of binding free energy of RNA with Rec domain of *Geo*Cas9 between WT, K267E and i*Geo*Cas9.

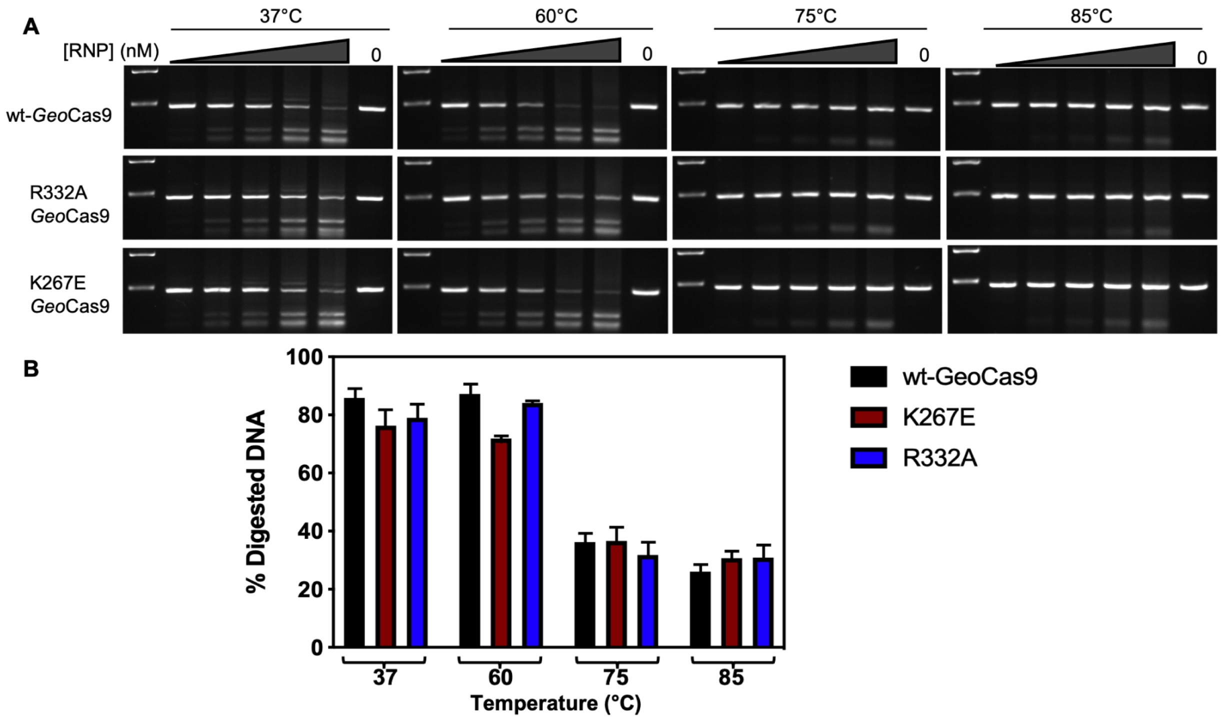


**Figure S17.** RNPs of varying concentrations of WT, K267E, or R332A *Geo*Cas9 and gRNA were incubated at 37, 60, 75, or 85 °C for 30 min, after which the RNPs were used for individual cleavage reactions. *In vitro* cleavage assays indicate no significant difference in temperature-dependent activity between WT, K267E, and R332A *Geo*Cas9. **(A)** RNP concentration in each lane (left-to-right) is 100, 200, 300, 600, or 900 nM and 0 nM (control) at each temperature tested. Molecular weight markers on agarose gels (top-to-bottom) are 600 and 400 basepairs. **(B)** Graph quantifying the DNA band intensity measurements on the agarose gel (A) using ImageJ. Data plotted as mean ± SD.

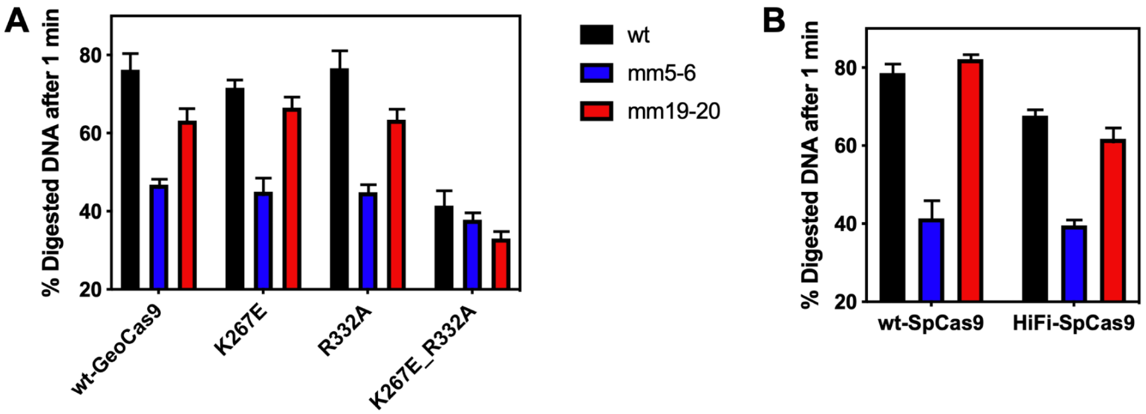


**Figure S18.** (**A**) Off-target *in vitro* cleavage assay with WT, K267E, R332A, and K267E/R332A *Geo*Cas9. (**B**) Off-target *in vitro* cleavage assay with WT and HiFi-*Spy*Cas9. Data plotted as mean ± SD. Legend: WT = on-target DNA substrate at the mouse Tnnt2 gene locus. mm5-6 = off-target DNA with the 5$^{th}$ and 6$^{th}$ nucleotide mismatches from the PAM seed site. mm19-20 = off-target DNA with the 19$^{th}$ and 20$^{th}$ nucleotide mismatches from the PAM seed site. Full DNA sequences of the substrates for the *Geo*Cas9 and *Spy*Cas9 can be found in Tables S2 and S3, respectively.

| Wild type | K267E | | R332A |
|---|---|---|---|
| $k_{ex}$ ($s^{-1}$) = 147 ± 41 | $k_{ex}$ ($s^{-1}$) = 376 ± 89 | | $k_{ex}$ ($s^{-1}$) = 142 ± 28 |
| F256 H-N | A249 H-N | V349 H-N | Q257 H-N |
| N266 H-N | I265 H-N | L359 H-N | G275 H-N |
| K267 H-N | L268 H-N | D362 H-N | A276 H-N |
| L270 H-N | L270 H-N | F363 H-N | R277 H-N |
| G275 H-N | I271 H-N | D376 H-N | L286 H-N |
| A276 H-N | S272 H-N | E388 H-N | T312 H-N |
| I297 H-N | G275 H-N | N399 H-N | Y319 H-N |
| D311 H-N | D281 H-N | N404 H-N | K326 H-N |
| Y313 H-N | L286 H-N | L406 H-N | N330 H-N |
| K315 H-N | K296 H-N | N412 H-N | F333 H-N |
| Y319 H-N | D310 H-N | L413 H-N | K344 H-N |
| K354 H-N | Y313 H-N | E434 H-N | V349 H-N |
| S356 H-N | Y319 H-N | V438 H-N | L359 H-N |
| Y387 H-N | E329 H-N | R445 H-N | D376 H-N |
| E409 H-N | N330 H-N | | M394 H-N |
| L413 H-N | F333 H-N | | D403 H-N |
| I429 H-N | E335 H-N | | N412 H-N |
| | R343 H-N | | L413 H-N |

**Table S1.** Residues fit to a global $k_{ex}$ in $^{1}H$-$^{15}N$ CPMG relaxation dispersion analysis of WT, K267E, and R332A *Geo*Rec2. A very small number of other resonances displaying curved CPMG profiles could not be globally fit and were excluded from this list. In all samples (*i.e.* WT, K267E, and R332A), the global fit was found to be the best statistical model.

| | |
|---|---|
| sgRNA | UUGCACCUACCUUCUGGAUGUACGUCAUAGUUCCCCUGAGAAAUCAGGGUUACUAUGAUAAGGGCUUUCUGCCUAA<br>GGCAGACUGACCCGCGGCGUUGGGGAUCGCCUGUCGCCCGCUUUUGGCGGGCAUUCCCCAUCCUU |
| On-Target DNA | CAAAGAGCTCCTCGTCCAGTGGGAAGAGAGCTGATCTCATTTGTAAGGAATACCCTCTTCATCCCCCACCCTTGCCATTG<br>ATCTATTCATTCCATCTCCATGACAACAGGAAGAGAGGGCCCGGCGTGAGGAGGAGGAGAACAGGAGGAAGGCTGAGG<br>ATGAGGCCCGGAAGAAGAAGGCTCTGTCCAACATGATGCACTTTGGAGGGTACATCCAGAAGGTAGGTGCAAAGCAGC<br>ATCGGGCACCAGGACACCCCAGTGTATCCTCAAGGCCGCCTTTGCTTGGATCCATGAAGAAATTCCCAACTGCTGTGGC<br>TGAAGTCTAAGGTCTGCTCATGTCTAGCCCCTGAGCTGTCTATCAGCCTGACCATGGCTTCAGTAGGAGGGCTCTGCTG<br>TGTGTGACAGTTAGAACACTAATATGTCTCCAAATTCTGGCTCCCCAAAGGGACAACTGGGAGAATCTTGGGTCCTGGA<br>GTCCAT |
| Off-Target DNA<br><br>PAM proximal-mismatch (5-6bp, AT>CA) | CAAAGAGCTCCTCGTCCAGTGGGAAGAGAGCTGATCTCATTTGTAAGGAATACCCTCTTCATCCCCCACCCTTGCCATTG<br>ATCTATTCATTCCATCTCCATGACAACAGGAAGAGAGGGCCCGGCGTGAGGAGGAGGAGAACAGGAGGAAGGCTGAGG<br>ATGAGGCCCGGAAGAAGAAGGCTCTGTCCAACATGATGCACTTTGGAGGGTACcaCCAGAAGGTAGGTGCAAAGCAGC<br>ATCGGGCACCAGGACACCCCAGTGTATCCTCAAGGCCGCCTTTGCTTGGATCCATGAAGAAATTCCCAACTGCTGTGGC<br>TGAAGTCTAAGGTCTGCTCATGTCTAGCCCCTGAGCTGTCTATCAGCCTGACCATGGCTTCAGTAGGAGGGCTCTGCTG<br>TGTGTGACAGTTAGAACACTAATATGTCTCCAAATTCTGGCTCCCCAAAGGGACAACTGGGAGAATCTTGGGTCCTGGA<br>GTCCAT |
| Off-Target DNA<br><br>PAM distal - mismatch(19-20bp, TG>CA) | CAAAGAGCTCCTCGTCCAGTGGGAAGAGAGCTGATCTCATTTGTAAGGAATACCCTCTTCATCCCCCACCCTTGCCATTG<br>ATCTATTCATTCCATCTCCATGACAACAGGAAGAGAGGGCCCGGCGTGAGGAGGAGGAGAACAGGAGGAAGGCTGAGG<br>ATGAGGCCCGGAAGAAGAAGGCTCTGTCCAACATGATGCACTTTGGAGGGTACATCCAGAAGGTAGGcaCAAAGCAGCA<br>TCGGGCACCAGGACACCCCAGTGTATCCTCAAGGCCGCCTTTGCTTGGATCCATGAAGAAATTCCCAACTGCTGTGGCT<br>GAAGTCTAAGGTCTGCTCATGTCTAGCCCCTGAGCTGTCTATCAGCCTGACCATGGCTTCAGTAGGAGGGCTCTGCTGT<br>GTGTGACAGTTAGAACACTAATATGTCTCCAAATTCTGGCTCCCCAAAGGGACAACTGGGAGAATCTTGGGTCCTGGAG<br>TCCAT |

**Table S2.** Nucleic acid sequences used in the *Geo*Cas9 *in vitro* off-target assay. The 23 base pair spacer sequence of gRNA is underlined. The spacer sequence within the DNA sequences is highlighted yellow, and the PAM are highlighted in blue.

| | |
|---|---|
| On-Target DNA | CAAAGAGCTCCTCGTCCAGTGGGAAGAGAGCTGATCTCATTTGTAAGGAATACCCTCTTCATCCCCCACCCTTGCCATTGATCTATTCATTCCATCTCCATG<br>ACAACAGGAAGAGAGGGCCCGGCGTGAGGAGGAGGAGAACAGGAGGAAGGCTGAGGATGAGGCCCGGAAGAAGAAGGCTCTGTCCAACATGATG<br>CACTTTGGAGGGTACATCCAGAAGGTAGGTGCAAAGCAGCATCGGGCACCAGGACACCCCAGTGTATCCTCAAGGCCGCCTTTGCTTGGATCCATGAA<br>GAAATTCCCAACTGCTGTGGCTGAAGTCTAAGGTCTGCTCATGTCTAGCCCCTGAGCTGTCTATCAGCCTGACCATGGCTTCAGTAGGAGGGCTCTGCTG<br>TGTGTGACAGTTAGAACACTAATATGTCTCCAAATTCTGGCTCCCCAAAGGGACAACTGGGAGAATCTTGGGTCCTGGAGTCCAT |
| Off-Target DNA<br><br>PAM proximal-mismatch (5-6bp, AA>CT) | CAAAGAGCTCCTCGTCCAGTGGGAAGAGAGCTGATCTCATTTGTAAGGAATACCCTCTTCATCCCCCACCCTTGCCATTGATCTATTCATTCCATCTCCATG<br>ACAACAGGAAGAGAGGGCCCGGCGTGAGGAGGAGGAGAACAGGAGGAAGGCTGAGGATGAGGCCCGGAAGctGAAGGCTCTGTCCAACATGATGC<br>ACTTTGGAGGGTACATCCAGAAGGTAGGTGCAAAGCAGCATCGGGCACCAGGACACCCCAGTGTATCCTCAAGGCCGCCTTTGCTTGGATCCATGAAG<br>AAATTCCCAACTGCTGTGGCTGAAGTCTAAGGTCTGCTCATGTCTAGCCCCTGAGCTGTCTATCAGCCTGACCATGGCTTCAGTAGGAGGGCTCTGCTGT<br>GTGTGACAGTTAGAACACTAATATGTCTCCAAATTCTGGCTCCCCAAAGGGACAACTGGGAGAATCTTGGGTCCTGGAGTCCAT |
| Off-Target DNA<br><br>PAM distal - mismatch(19-20bp, CA>AG) | CAAAGAGCTCCTCGTCCAGTGGGAAGAGAGCTGATCTCATTTGTAAGGAATACCCTCTTCATCCCCCACCCTTGCCATTGATCTATTCATTCCATCTCCATG<br>ACAACAGGAAGAGAGGGCCCGGCGTGAGGAGGAGGAGAACAGGAGGAAGGCTGAGGATGAGGCCCGGAAGAAGAAGGCTCTGTCagACATGATG<br>CACTTTGGAGGGTACATCCAGAAGGTAGGTGCAAAGCAGCATCGGGCACCAGGACACCCCAGTGTATCCTCAAGGCCGCCTTTGCTTGGATCCATGAA<br>GAAATTCCCAACTGCTGTGGCTGAAGTCTAAGGTCTGCTCATGTCTAGCCCCTGAGCTGTCTATCAGCCTGACCATGGCTTCAGTAGGAGGGCTCTGCTG<br>TGTGTGACAGTTAGAACACTAATATGTCTCCAAATTCTGGCTCCCCAAAGGGACAACTGGGAGAATCTTGGGTCCTGGAGTCCAT |

**Table S3.** DNA sequences used in the *Sp*Cas9 *in vitro* off-target assay. Sites of mismatched DNA are highlighted in red.

| | |
|---|---|
| Tnnt2 gRNA | UUGCACCUACCUUCUGGAUGUACGUCAUAGUUCCCCUGAGAAAUCAGGGUUACUAUGAUAAGGGCUUUCUGCC<br>UAAGGCAGACUGACCCGCGGCGUUGGGGAUCGCCUGUCGCCCGCUUUUGGCGGGCAUUCCCCAUCCUU |
| 8UZA gRNA | CACUGCAUUCUAGUUGUGGUUGUCAUAGUUCCCCUGAGAAAUCAGGGUUACUAUGAUAAG<br>GGCUUUCUGCCUAAGGCAGACUGACCCGCGGCGUUGGGGAUCGCCUGUCGCCCGCUUUUG<br>GCGGGCAUUCCCCAUCCUU |

**Table S4.** Guide RNA sequences used in *Geo*Cas9 MST measurements. The 39-nucleotide sequence of RNA used in MST and NMR studies of isolated *Geo*Rec is underlined.
